# ImmuneFM: Pre-training Foundation Model from Cytometry Data for Immunology Research

**DOI:** 10.1101/2025.07.09.664020

**Authors:** Sirui Ding, Sanchita Bhattacharya, Atul J. Butte

**Affiliations:** Bakar Computational Health Sciences Institute, University of California, San Francisco, CA, USA

## Abstract

Immunology is an essential field in the biomedicine domain, which plays an important role in oncology, vaccines, infection, etc. With the increasing amount of data available in immunology and artificial intelligence technique development, there is a need to develop data-driven AI methods in the field. However, the data from various immunology studies is very hard to integrate into an AI-ready dataset due to the lack of a standard. Moreover, independent immunology studies’ data lacks enough labels to train the supervised model. Motivated by these challenges, we curated a large-scale AI-ready cytometry dataset for immunology from the publicly available ImmPort portal. We design the framework to pre-train a foundation model, ImmuneFM, on the cytometry dataset. ImmuneFM can be applied to a wide range of downstream immunology diseases with fine-tuning on a limited number of labeled samples. The experiment results on eight downstream tasks demonstrate the superior performance of ImmuneFM compared to baseline deep learning and traditional methods.

## Introduction

Immunology is a foundational discipline in the biomedicine domain^1^, playing a critical role in advancing our understanding and treatment of diverse diseases - from infectious diseases^2^ to autoimmune disorders^3^ and oncology^4^. In recent years, transformative breakthroughs have emerged in this field. For example, mRNA COVID-19 vaccines demonstrate quick immune-based strategies in responding to the global pandemic^5^. Immunology also revolutionizes cancer treatment, e.g., the CAR-T cell therapy, which engineers the patient’s T cell to target cancer cells^6^. The immune system’s complexity, with its intricate network of cells, signaling pathways, and molecular interactions, necessitates advanced tools for comprehensive analysis. Immunology research generates vast volumes of data from observational and clinical trial studies. The ImmPort data portal^7^, a publicly accessible repository for immunology-related studies, currently hosts 1,256 studies spanning 176 diseases and includes over 7.5 million experimental results (as of April 2025). This explosion of immunological data presents a timely opportunity to leverage artificial intelligence (AI) for both clinical diagnostics and scientific discovery.

Recent advancements in AI have shown great promise in decoding the complexities in immunology, offering new insights into immune responses and potential therapeutic targets^8^. For instance, AI can be used to discover new biomarkers for immune oncology^9^; AI can also be applied in drug discovery and drug repurposing^10,11^. However, the application of AI in immunology still faces significant challenges, particularly when dealing with independent studies involving very few subjects. One of the primary challenges in applying AI to immunology is the scarcity of large, well-curated datasets and the lack of standardization of immunology data. Unlike other fields where massive, labeled datasets are readily available, immunological studies often involve small patient cohorts due to the rarity of certain conditions, ethical considerations, and the high cost of data collection and labeling^12^. This data scarcity limits the ability of traditional AI models to learn robust patterns and make accurate predictions^13^. The other challenge is the heterogeneity of immune responses, which makes it hard for a supervised training AI model to be adapted to various types of diseases^14^. So, these two challenges make it a dilemma that immunologists need to train an AI model for each study, but each study lacks enough labeled data. These challenges highlight the need for a new paradigm in how AI is applied to immunology^15^, moving from models trained on isolated, small datasets to those that can integrate and learn from diverse sources of immunological data.

With the rapidly increasing data accessibility in the public domain and promising performance of self-supervised learning, the foundation model becomes the new paradigm in the biomedical domain^16^. A foundation model is a general AI model pre-trained on large-scale and wide-range datasets, which can support various downstream tasks. Large language model (LLM) is one type of foundation model^17^. For example, a biomedicine foundation model can be used for disease diagnosis, clinical decision support, scientific discovery, etc^18^. Driven by big data and large-scale models, foundation models have shown strong power in the biomedicine domain. For example, Steinberg et al. designed MOTOR, a foundation model pre-trained on structured EHR data^19^, Wang et al. pre-trained a pathology foundation model which can be applied for cancer studies^20^, Yang et al. pre-trained GatorTron, a large language model on clinical text^21^. In the field of immunology, Kim et al.^22^ pre-trained a masked autoencoder on the cytometry data specifically for COVID-19 analysis. However, there remains a lack of a generalizable immunology foundation model capable of addressing a broader spectrum of diseases and conditions, such as oncology, transplantation, and autoimmune disorders.

In this work, we introduce **ImmuneFM**, a foundation model pre-trained on large-scale cytometry data from 53 immunology studies in the ImmPort repository, spanning eight domains including vaccine response, autoimmune disease, oncology, transplantation, infection, allergy, preterm birth, and general immune response. With cytometry accounting for nearly 70% of ImmPort results, we curated a dataset of over 100 million single cells and 203 commonly used panel markers. ImmuneFM follows a two-stage framework: during pre-training, raw cytometry data are normalized and masked for self-supervised learning using a Transformer model; during fine-tuning, cell embeddings are aggregated and passed to a multi-layer perceptron (MLP) for classification. We evaluated ImmuneFM on eight downstream tasks, such as liver cancer staging^23^, allergy prediction^24^, autoimmune disease classification^25^, and HIV/COVID diagnosis^26–28^, where it consistently outperformed baseline models. Further interpretability analysis at the cell and marker levels enabled both validation of known findings and discovery of new insights across three case studies, demonstrating ImmuneFM’s potential for both clinical application and scientific discovery.

In summary, we have three-fold contributions in this study:

- We curated a large-scale AI-ready cytometry dataset covering a wide range of focusing immunology areas, from oncology to transplant.
- We propose and pre-train ImmuneFM, a foundation model for the immunology field to support clinical diagnosis and scientific discovery.
- We evaluate the ImmuneFM on 8 downstream tasks of different diseases, which demonstrate the superior accuracy and interpretability of the model.

## Methods

### Data preparation

#### Pre-training Dataset curation

We built the cytometry dataset from the ImmPort platform, which is an open-source database for immunology studies. As shown in Figure 2, we integrate the cytometry data collected from 53 different studies covering 8 focusing areas on immunology, including vaccine, COVID-19, oncology, etc. Each immunology study used different panel markers in the cytometry assay, so we manually created a commonly used panel marker set, which contains 203 frequently used markers, e.g., CD4, CD8, etc. Due to the extremely high volume of cytometry data in the 53 studies, we sample 10% of the single cells from each subject in the study. The final curated cytometry dataset is shaped in a matrix, where each row represents a single cell, and each column represents a panel marker. The detailed marker name and study ID of the pre-training dataset are presented in Figure. 2.

**Figure 1.**
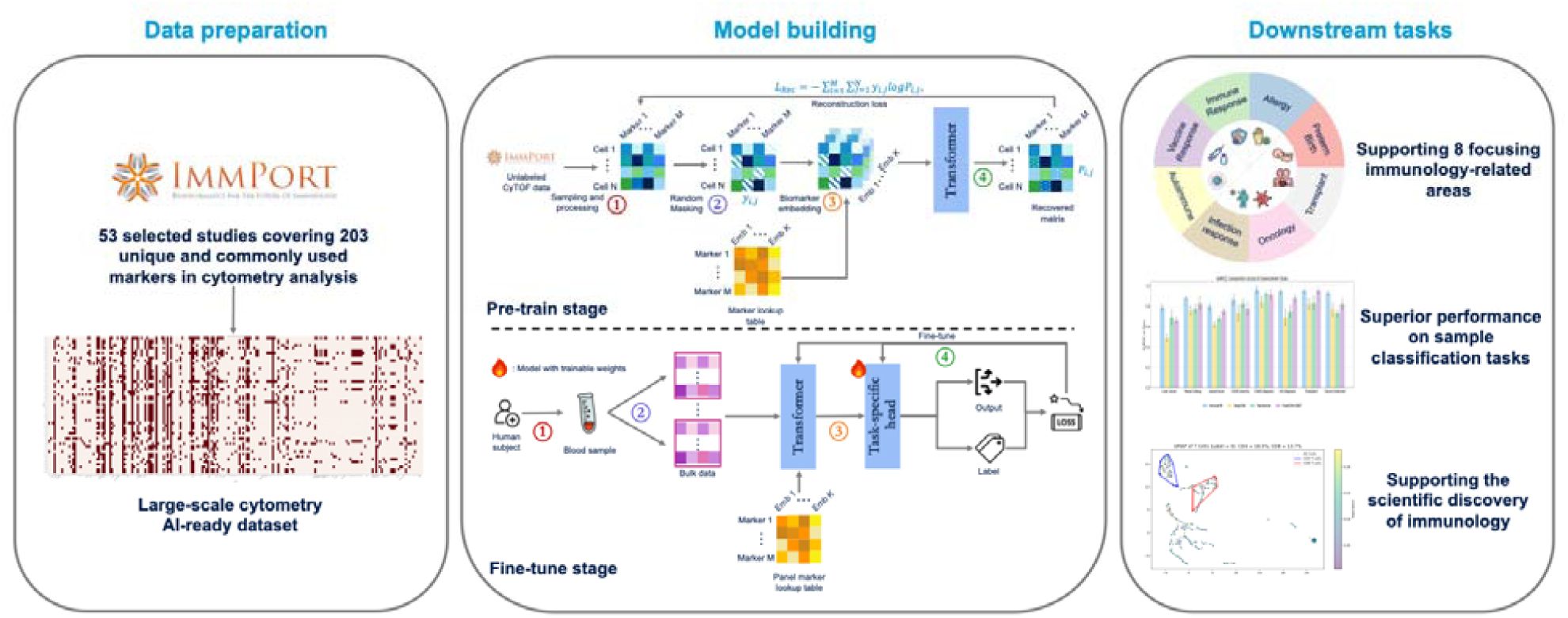
Workflow overview of ImmuneFM. 53 public studies selected from the ImmPort portal were used to integrate a large-scale AI-ready cytometry dataset. ImmuneFM was pre-trained on this dataset and fine-tuned for downstream sample classification tasks.

**Figure 2.**
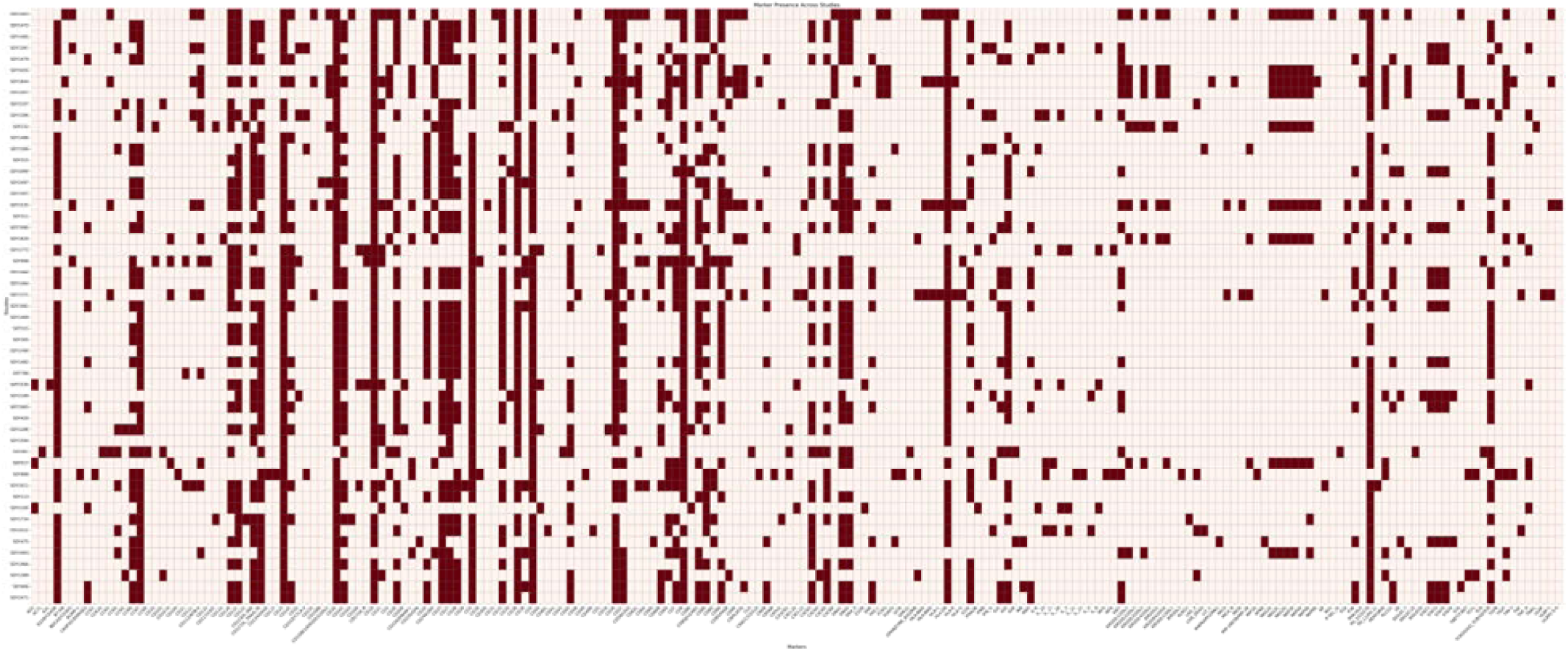
Study and marker details of the curated cytometry dataset for pre-training. There are 53 studies from ImmPort that are included, covering 203 unique panel markers.

#### Pre-processing

Because the studies are conducted by different labs on different samples, we apply the widely used normalization function to transform the single-cell data in cytometry^29^. Because we need to embed each marker, which requires the marker value to be an integer, we linearly scale the expression value with MinMaxScaler and then bin the values into discrete integers inspired by NLP techniques^30^ and gene expression binning^31^.

### The ImmuneFM model

#### Transformer backbone

We built the immunology foundation model, the ImmuneFM, with the Transformer backbone. The input data is a cytometry matrix that can be embedded like the word token embedding in the BERT model. Meanwhile, the order of marker columns doesn’t influence the semantic meaning of the cytometry matrix. Thus, we don’t need a positional embedding like an NLP task. Because the cytometry marker number is not like the gene number, which can be nearly 20,000^31^, the commonly used Transformer^32^ can achieve competitive performance with a 203-panel marker length. The core formulas for multi-head self-attention are as follows:

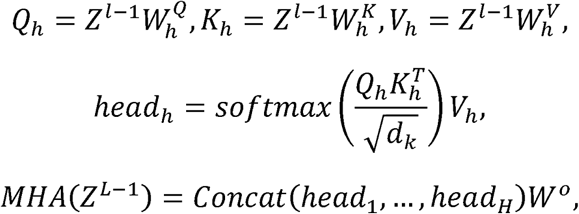

where Q,J,V are the query, key, and value vectors. The ,,wi is a single attention

***Panel marker embedding.*** Like the word embedding to process text^33^, we are also inspired to embed each marker with a learnable representation. Because the embedding can ensure that similar markers have close and similar representations. We employ a trainable embedding layer to encode each panel marker with a look-up table. We can learn the inter-marker relations with this embedding, and we name this layer “Marker2Vec” as follows.

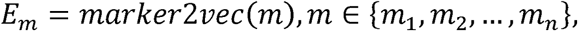

where each panel marker m is mapped to a vector cm via mw’k,’2,;,c.

***Marker expression embedding.*** The value of each panel marker reflects the expression levels of specific proteins or molecules. The marker expression can be considered as the existing occurrence of different biomarkers from a single cell. Inspired by this, we follow the conventional method to convert the continuous value of a marker expression into a discrete integer value by binning it into an *N*-dimensional vector that can be input into ImmuneFM as follows.

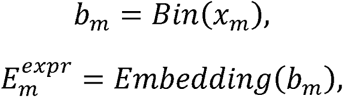

where xm is the expression value of marker m.

**Sample/bulk level representation.** In the cytometry analysis, the sample or bulk classification is the focusing task, which is usually used for disease diagnosis, treatment effectiveness analysis, etc. A sample contains a bunch of single cells, e.g., 10k cells for each sample. We will encode each cell with the pre-trained backbone and aggregate the embedding into a sample-level tensor with mean pooling as follows.

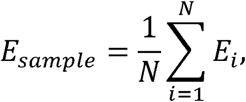

where N is the number of cells in a sample.

**Sample classification head.** Then, a task-specific head, e.g., several linear layers, will be applied after the Transformer backbone to process the sample-level tensor and output the classification results. The classification head can be represented as follows.

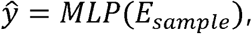

where the y’ is the model output from a multi-layer perceptron ("LPo.

#### Pre-train and fine-tune

***Self-supervised learning with masking.*** In this work, we use the masking- regeneration task to pre-train the ImmuneFM. We will randomly mask the non-zero items in the input cytometry matrix and reconstruct the masked items with the remaining values in the matrix from the Transformer backbone output. Cross-entropy is used as the loss function to optimize as follows:

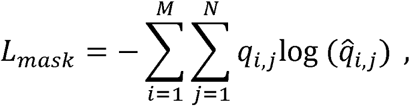

where N is each sample’s cell number, " is the number of markers, which is 203 in the and 1th cell) in the original cytometry matrix and reconstructed output from the

***Supervised fine-tuning on downstream tasks.*** After pre-training the backbone Transformer, we will add a task-specific head to process the outputs. This task-specific head can be several layers of an MLP. In this work, we apply the MLP to output the classification results. We used cross-entropy to optimize the ImmuneFM as follows.

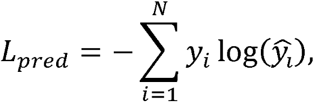

where yi,y∼i indicate the ground truth and output from the model.

#### Model interpretation

We use the gradient-based class activation mapping (Grad-CAM) to calculate the importance of each cell. To interpret the model’s prediction, we compute the gradient of the predicted class score with respect to each cell embedding. These gradients are then combined with the original embeddings to produce importance scores for individual cells. This highlights which cells contributed most to the model’s decision, providing biological insights at the single-cell level. The interpretation calculation process can be represented as follows.

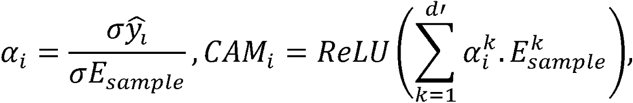

where vi computes the gradient of class y∼i w.r.t. each cell embedding csample . The importance score i for each cell is calculated by multiplying the gradient the the embedding.

#### Baseline methods

***FlowSOM + GBDT.*** We apply FlowSOM to cluster the cells in a sample. And then calculate statistical features from these clusters, including cluster percentage in the sample, mean fluorescence intensity (MFI) of panel markers. Then we used these derived features to train and test a GBDT classifier.^34^

***CNN.*** We compare with the CNN model specifically designed for cytometry sample classification, which is proposed by Hu et al.^29^. We follow the hyperparameter settings as described in the manuscript.

***Transformer.*** We compare with the Transformer backbone model without pre-training on the large-scale cytometry dataset. The hyperparameter settings for the plain Transformer model are the same as ImmuneFM.

#### Implementation details

**For the pre-training of ImmuneFM**, the training epoch is set to 100, and the learning rate is 1e-4. The batch size for pre-training is 1024 cells. The train-validation split is 0.95/0.05. The Transformer’s dimensions, depth, and heads are 128, 6, and 8, respectively. The bin number is 10. The masking ratio is 0.25. We use the Adam optimizer to pre-train the ImmuneFM.

**For the fine-tuning of ImmuneFM,** the epoch is set to 50, and the learning rate is 1e-4 with the Adam optimizer. The cell number in each sample is set to 512, 256, and 128. The batch size for pre-training is 4.

**For the baseline models,** the DeepCNN model follows the implementation and hyperparameter settings of Hu et al.’s work^29^. The Transformer model is the same architecture as ImmuneFM. FlowSOM is implemented with the Python package by Couckuyt et al^34^. The GBDT model is implemented by the Scikit Learn Python package.

All the models are implemented with Python and packages including PyTorch, Scikit Learn, etc. The training and testing of the model are conducted on the UCSF Wynton platform.

#### Downstream task curation

We evaluated the ImmuneFM across eight downstream tasks as shown in Table 1 spanning various human immunology diseases, including a cross-species validation using a mouse dataset. These tasks were derived from the publicly available datasets in ImmPort, including SDY1733 (liver cancer), SDY2015(peanut allergy), SDY997(lupus nephritis), SDY1708(COVID-19), SDY2011 (COVID-19), SDY1535 (HIV), SDY788

**Table 1.**
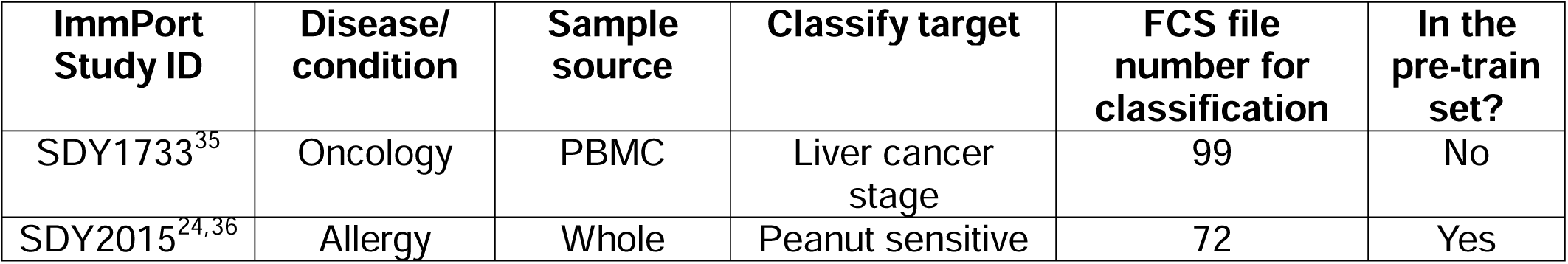
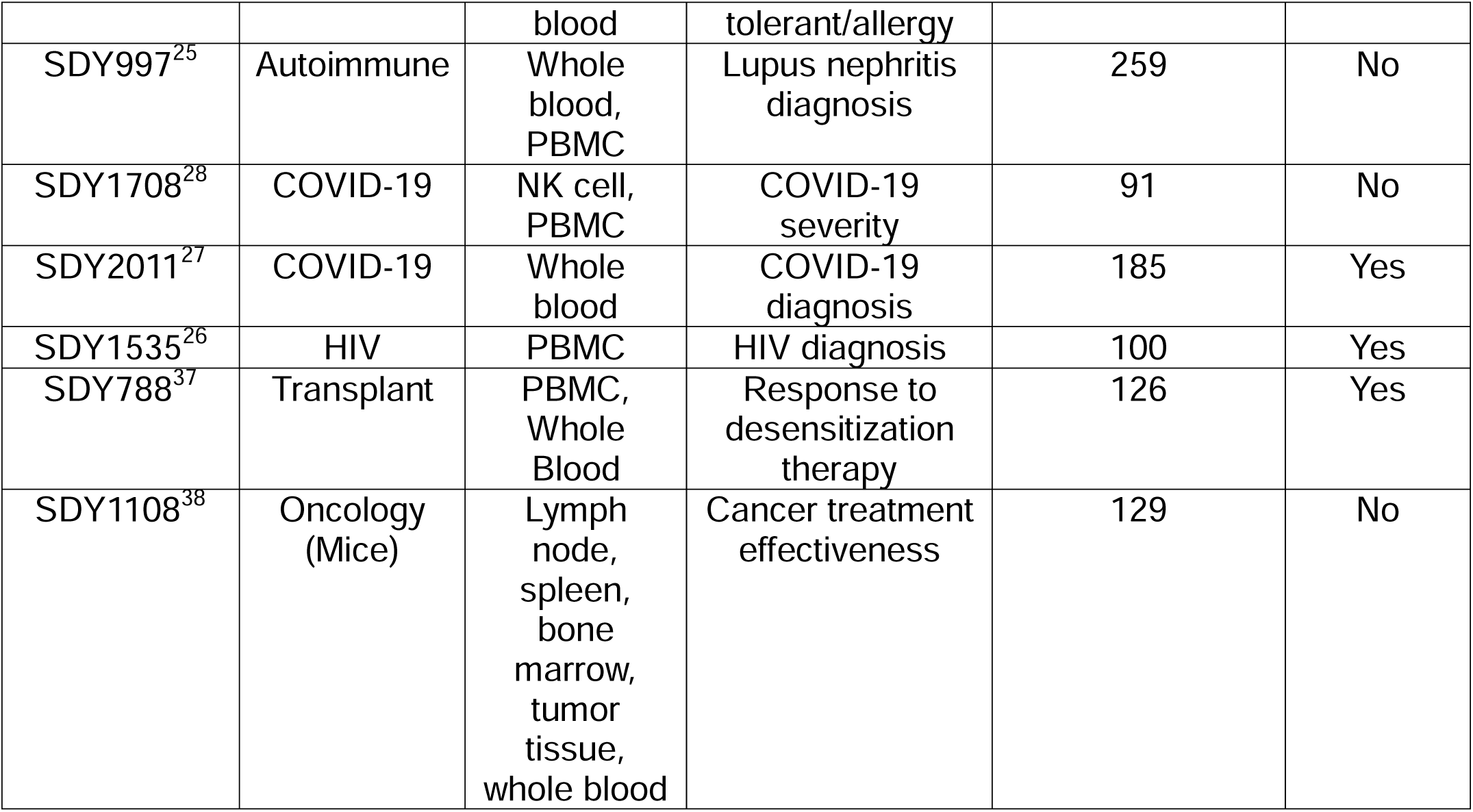
Statistical characteristics of downstream task studies.

(kidney transplant), and SDY1108 (cancer).

#### Evaluation metrics

For the binary classification task, we use the area under the receiver operating characteristic curve (AUROC) as the evaluation metric. For the multi-class classification task, we use balanced accuracy (Bacc) as the evaluation metric. The formulas to calculate AUROC and Bacc are as follows.

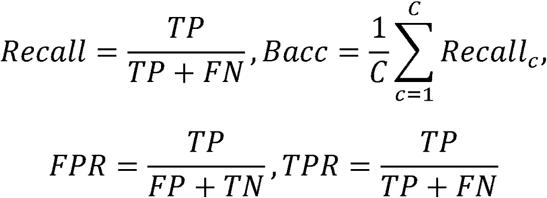

#### Benchmarking against state-of-the-art methods

We benchmark ImmuneFM on eight downstream tasks against state-of-the-art baseline methods and under different hyperparameter settings (See Methods). We have several observations and insights from the results in Figure 3 (a), (b), and (c).

**Figure 3.**
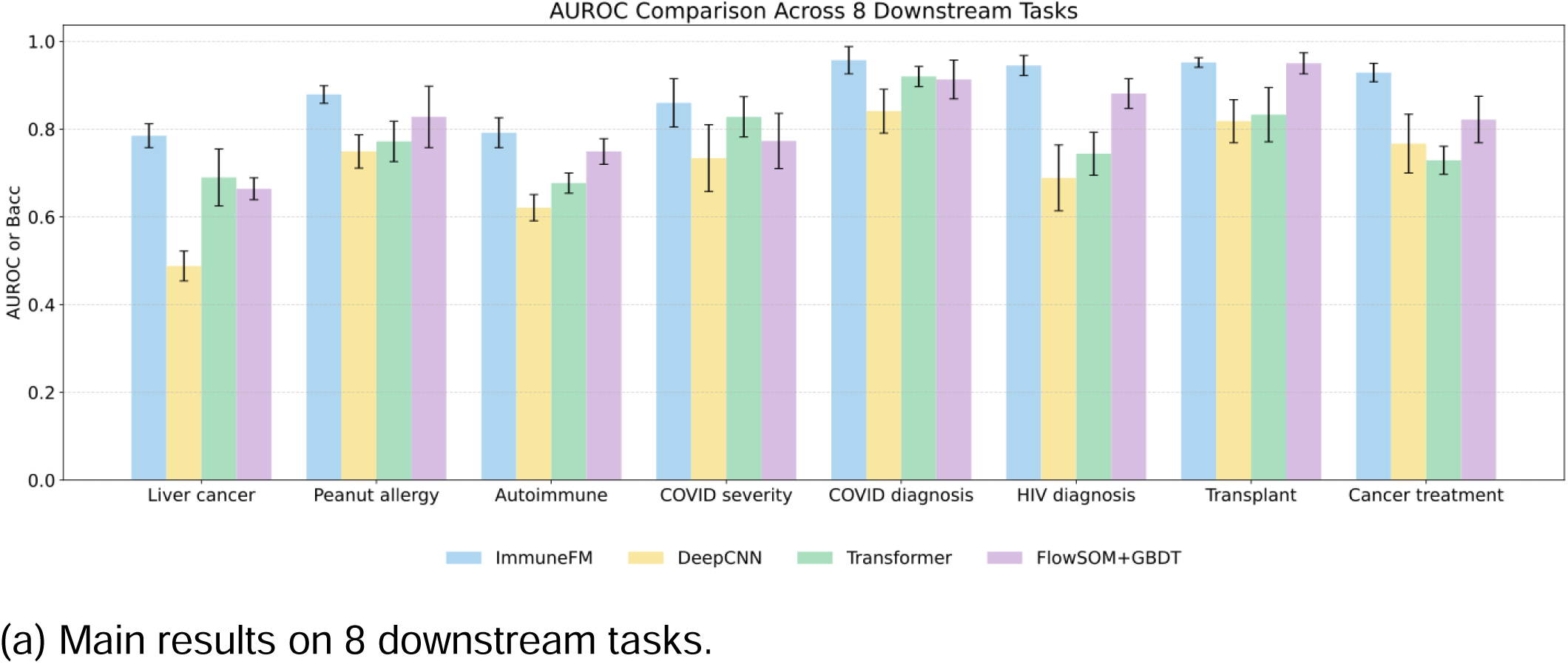
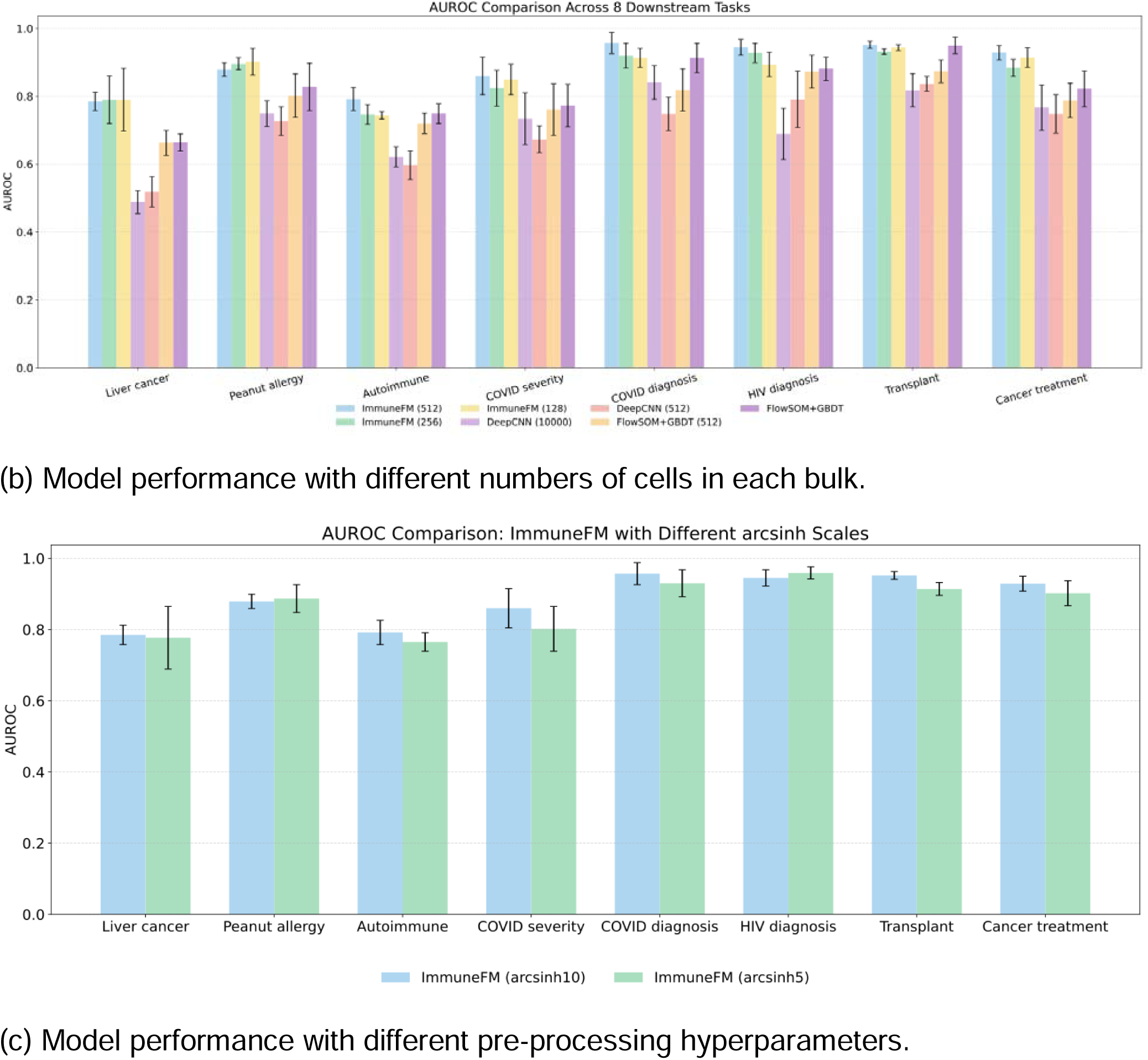
Sample classification performance on 8 different immunology diseases and conditions. For binary classification, AUROC is the metric for evaluation. For multi-class classification, balanced accuracy (Bacc) is the metric for evaluation.

As shown in Figure 3 (a), we found that ImmuneFM significantly outperforms DeepCNN^29^, Transformer^31^, and FlowSOM+GBDT^34^ across the downstream sample classification tasks. Firstly, we observed that the CNN model has the lowest performance, which indicates the drawback of the CNN model under the limited number of labeled samples. Secondly, the Transformer without pre-training outperforms the CNN model, which implies a more suitable model architecture in cytometry classifications. Compared to the pre-trained ImmuneFM, the plain Transformer model has significantly worse performance, indicating the necessity and boosting effect of pre-training. Thirdly, the traditional method, such as FlowSOM+GBDT, achieved surprisingly competitive performance. The traditional method with feature engineering, like FlowSOM, outperforms the deep learning models, including CNN and Transformer. It makes sense especially under the circumstances of limited labeled samples. The traditional method with feature engineering needs fewer labeled samples than supervised deep learning.

As shown in Figure 3 (b), the insight is that the cell number in each sample/bulk didn’t have a significant effect on the performance of deep learning models. We can observe that the ImmuneFM with much fewer cells, e.g., 256 or 128 cells in each bulk, also has competitive performance compared to 512 cells as input. This phenomenon is also observed in the CNN model. However, the cell number in each bulk has a larger effect on the FlowSOM+GBDT. We can observe that the performance of FlowSOM+GBDT drops with fewer cells in each sample. This is mainly due to the traditional method relies heavily on feature engineering which is sensitive to the number of cells. As shown in Figure 3 (c), we can find that ImmuneFM is robust with different pre-processing hyperparameters, e.g., the arcsinh factor. This result demonstrates the robustness of the sensitivity of ImmuneFM to the hyperparameters.

#### Interpretable analysis for scientific results

To further understand how ImmuneFM makes predictions and to explore its capacity for scientific insight, we conducted an interpretable analysis at both the marker and single-cell resolution. Building on the strong classification performance, this analysis aimed to explain model decisions and uncover biologically meaningful patterns. We focused on three case studies—COVID-19, HIV, and liver cancer—and used class activation mapping to visualize the importance of individual cells and markers in the cytometry input matrix (Figure 4). At the marker level, we computed the average contribution of each marker across positive and negative cohorts and compared expression levels to highlight differentially informative biomarkers (Figure 5). At the cell level, we leveraged these key markers to identify immune cell populations and assess their relevance to sample classification (Figure 6). This dual-level interpretation not only aligned with findings from the literature but also revealed novel cellular and molecular features, underscoring ImmuneFM’s potential for enabling data-driven discovery in immunology. We provided clinical and scientific insights based on the marker and cell interpretation results. These insights are either validated with the previous literature with scientific evidence or new discoveries that haven’t been widely reported in previous works.

**Figure 4.**
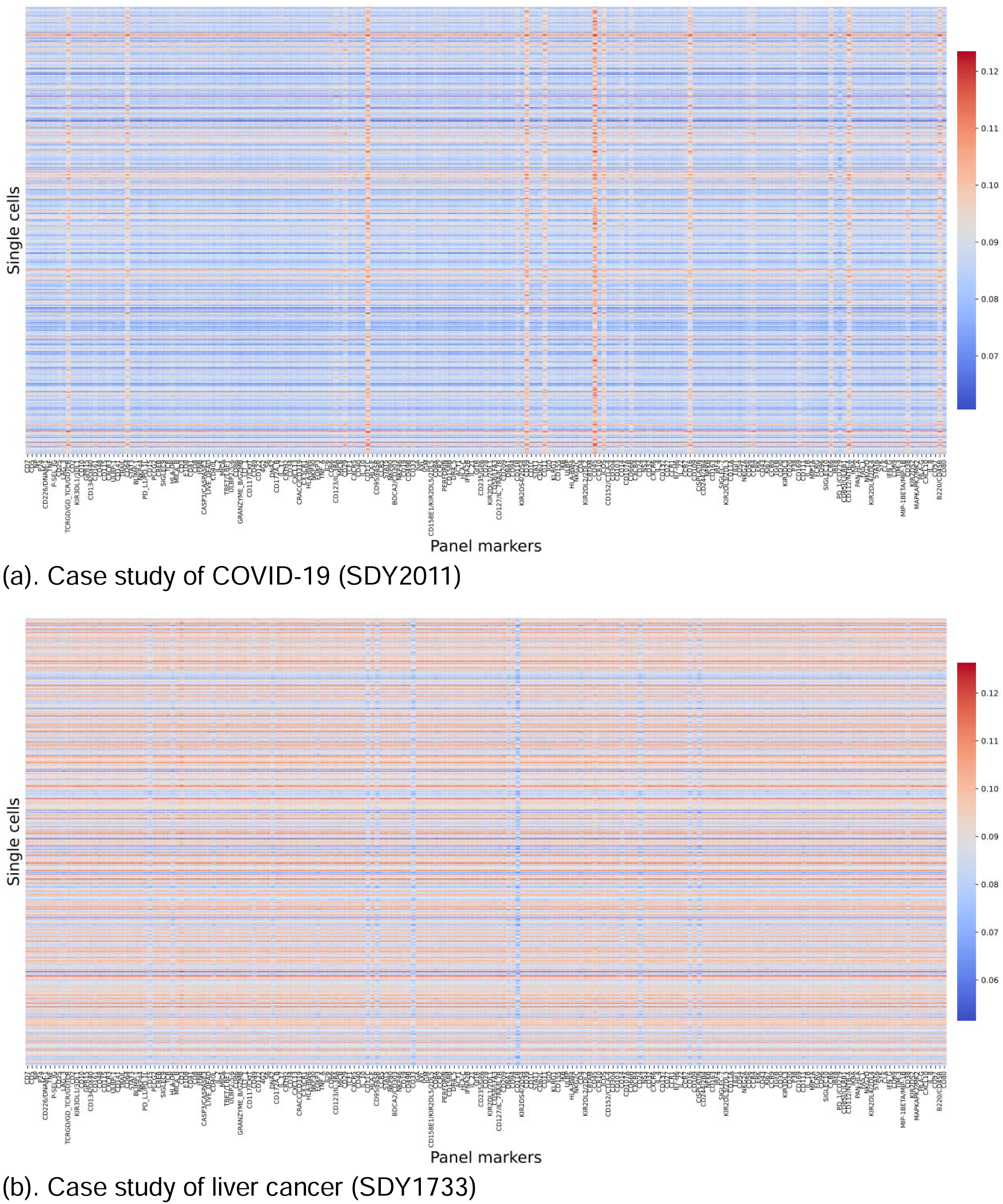
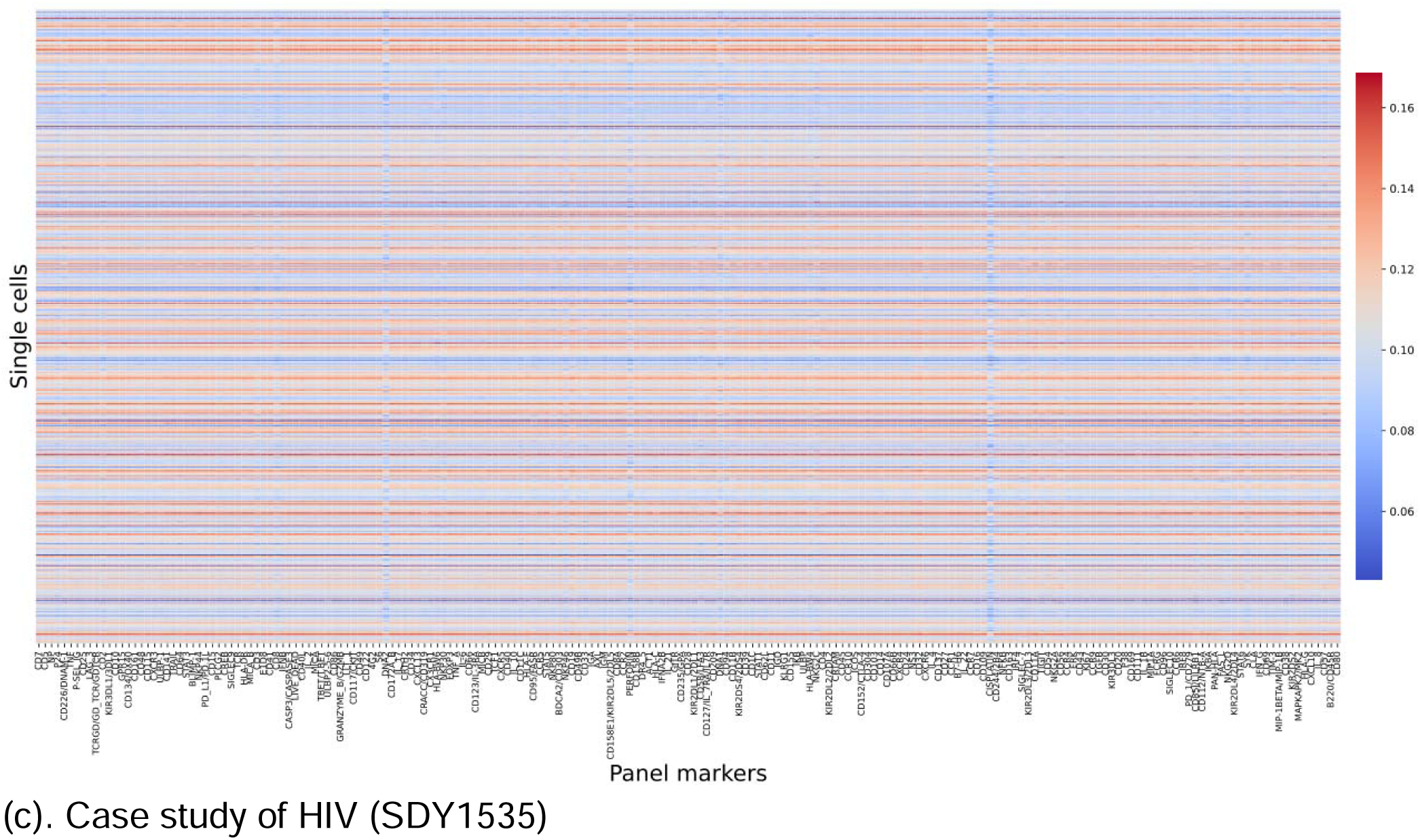
Heatmap on input cytometry matrix. Each row represents a single cell, and each column represents a unique panel marker. Color indicates the importance of each cell or marker.

**Figure 5.**
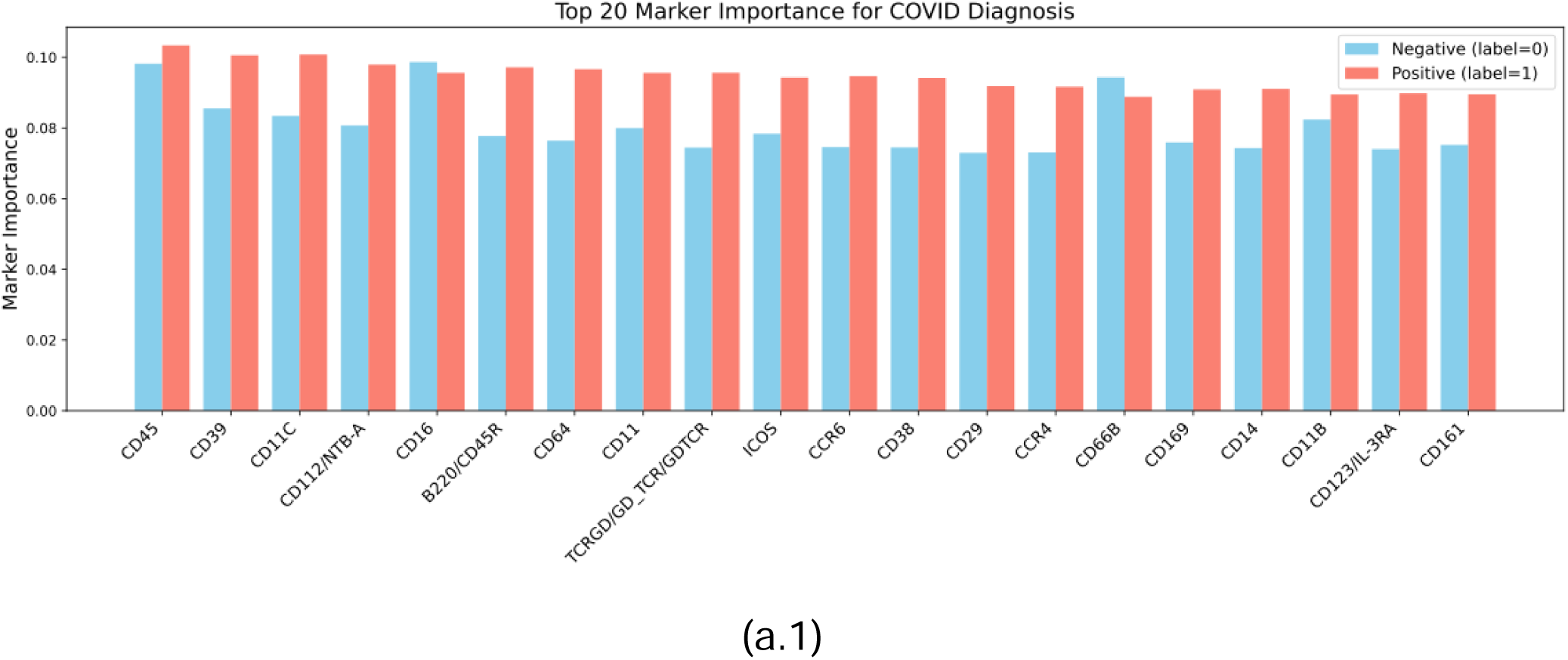
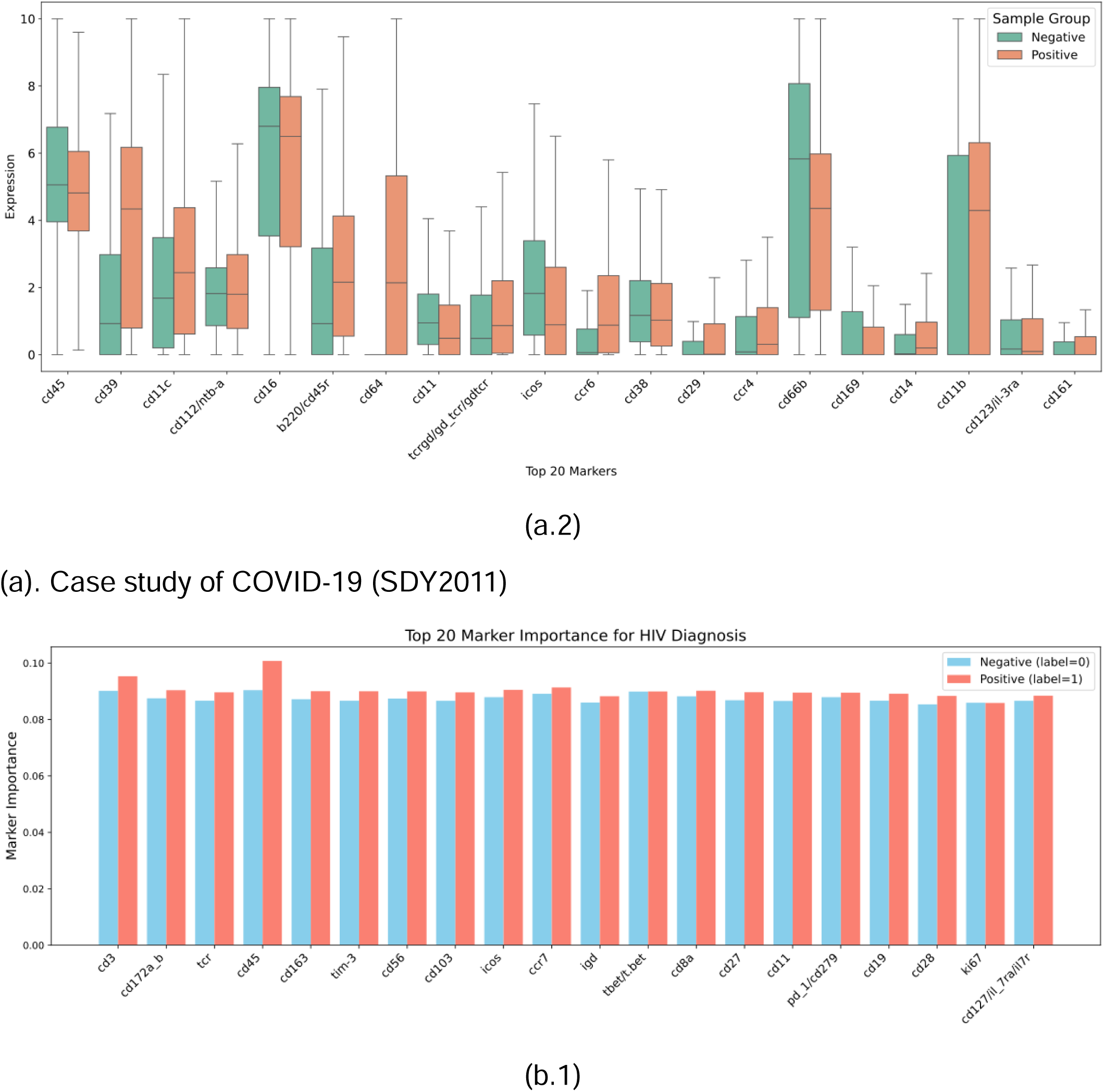
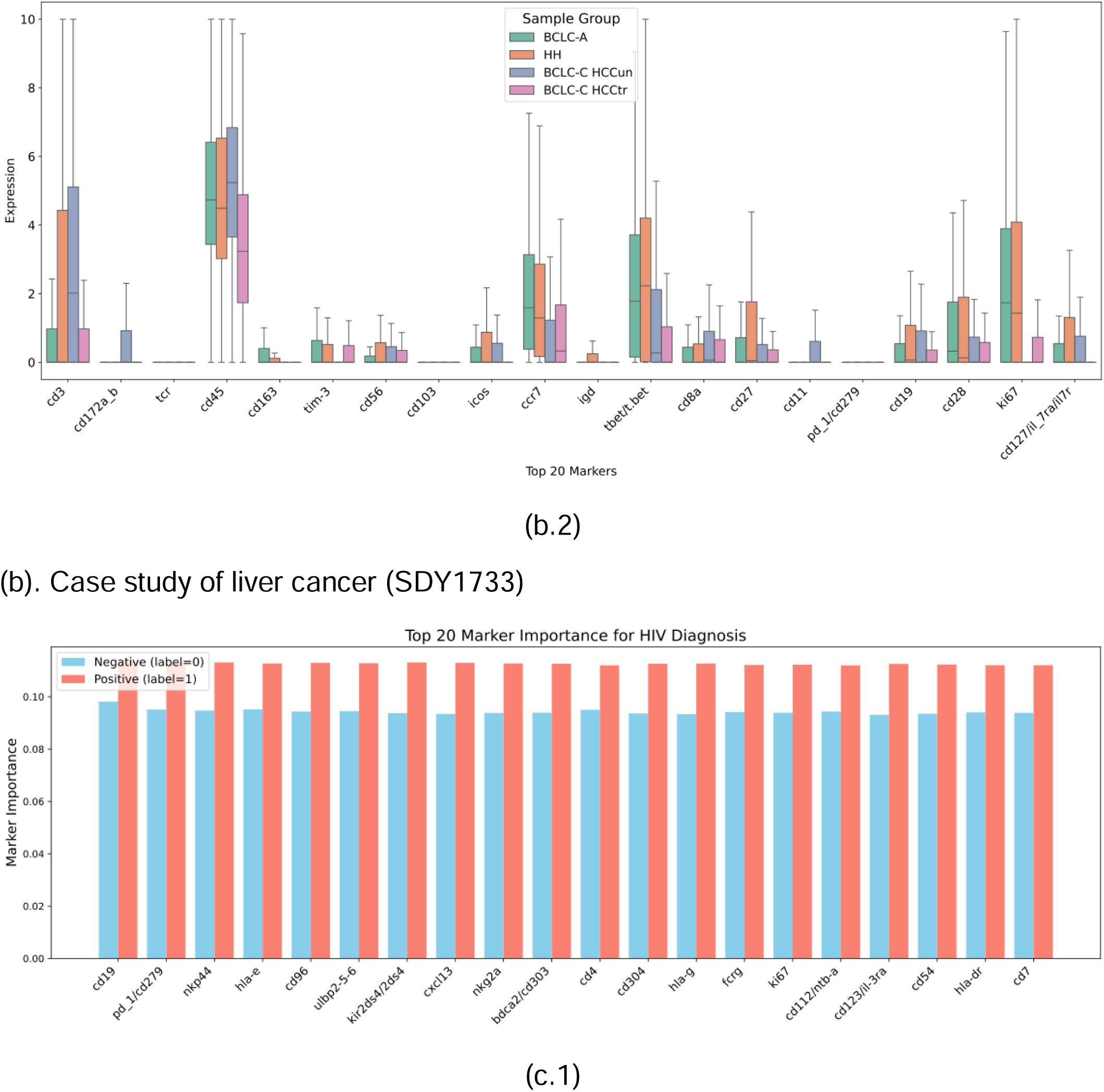
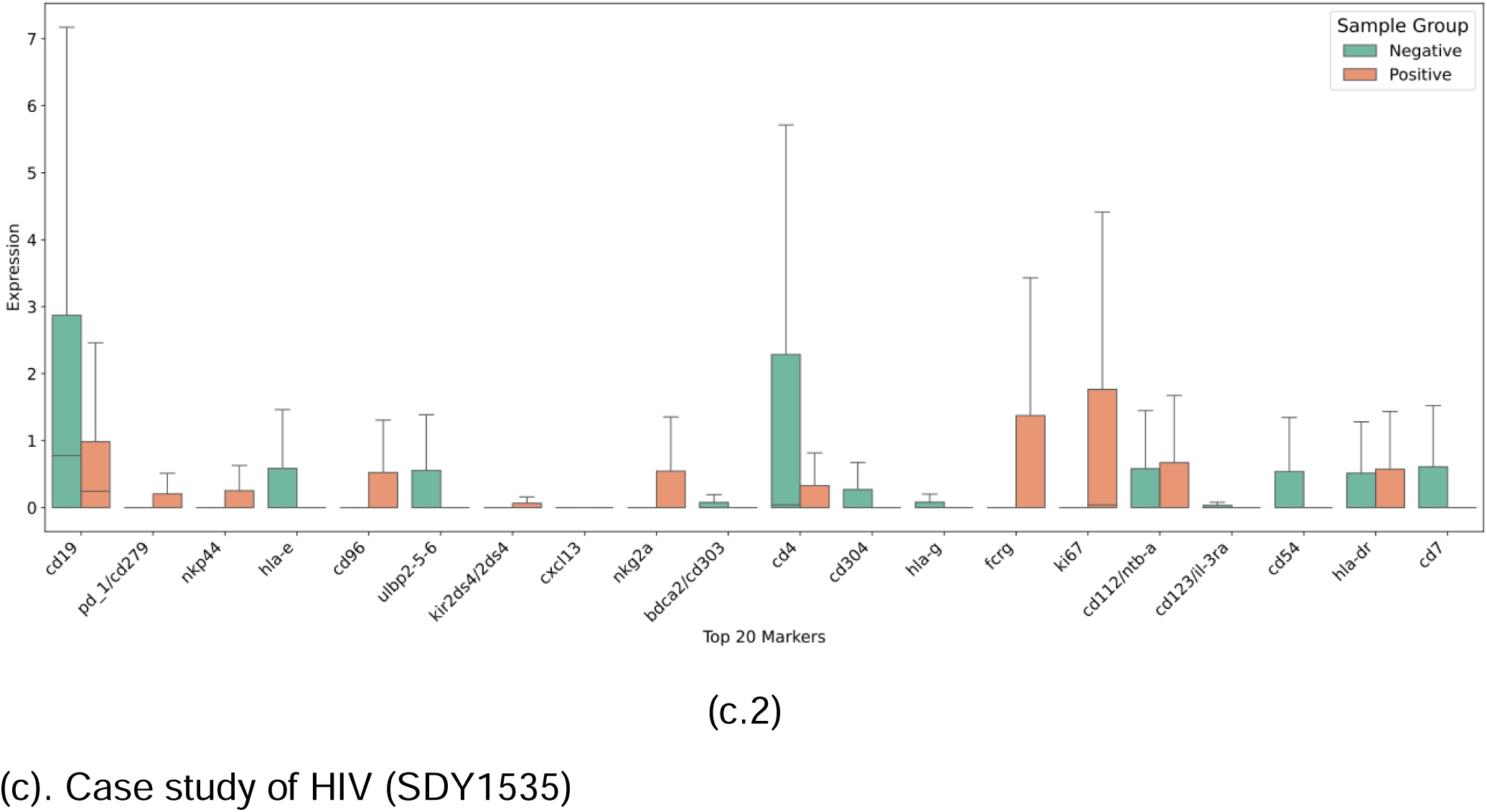
Importance and expression of panel markers in positive and negative patient groups. The subfigure (1) indicates the importance of the marker that contributes to the classification results, and subfigure (2) shows the expression level of important markers in different subgroups.

**Figure 6.**
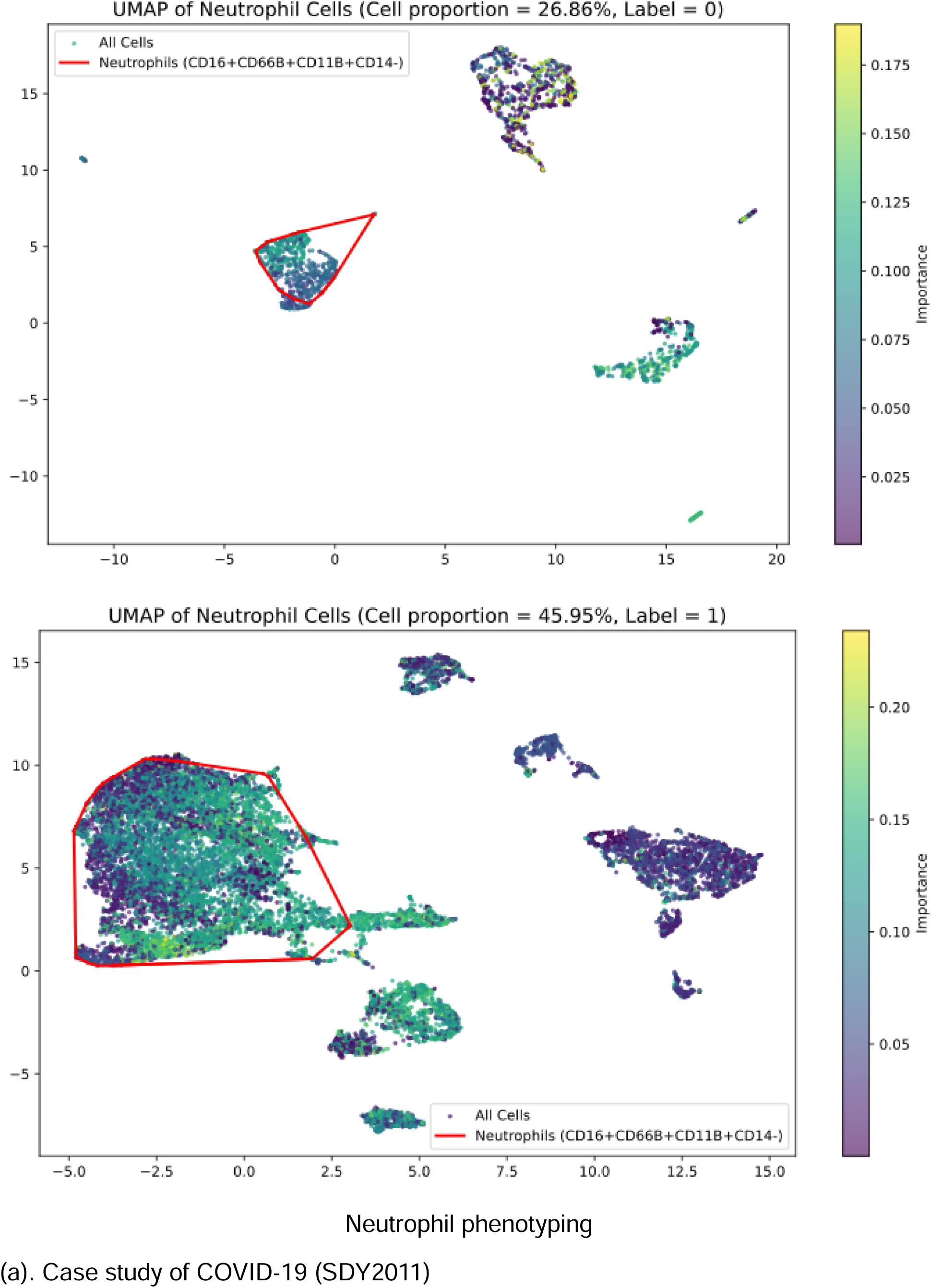
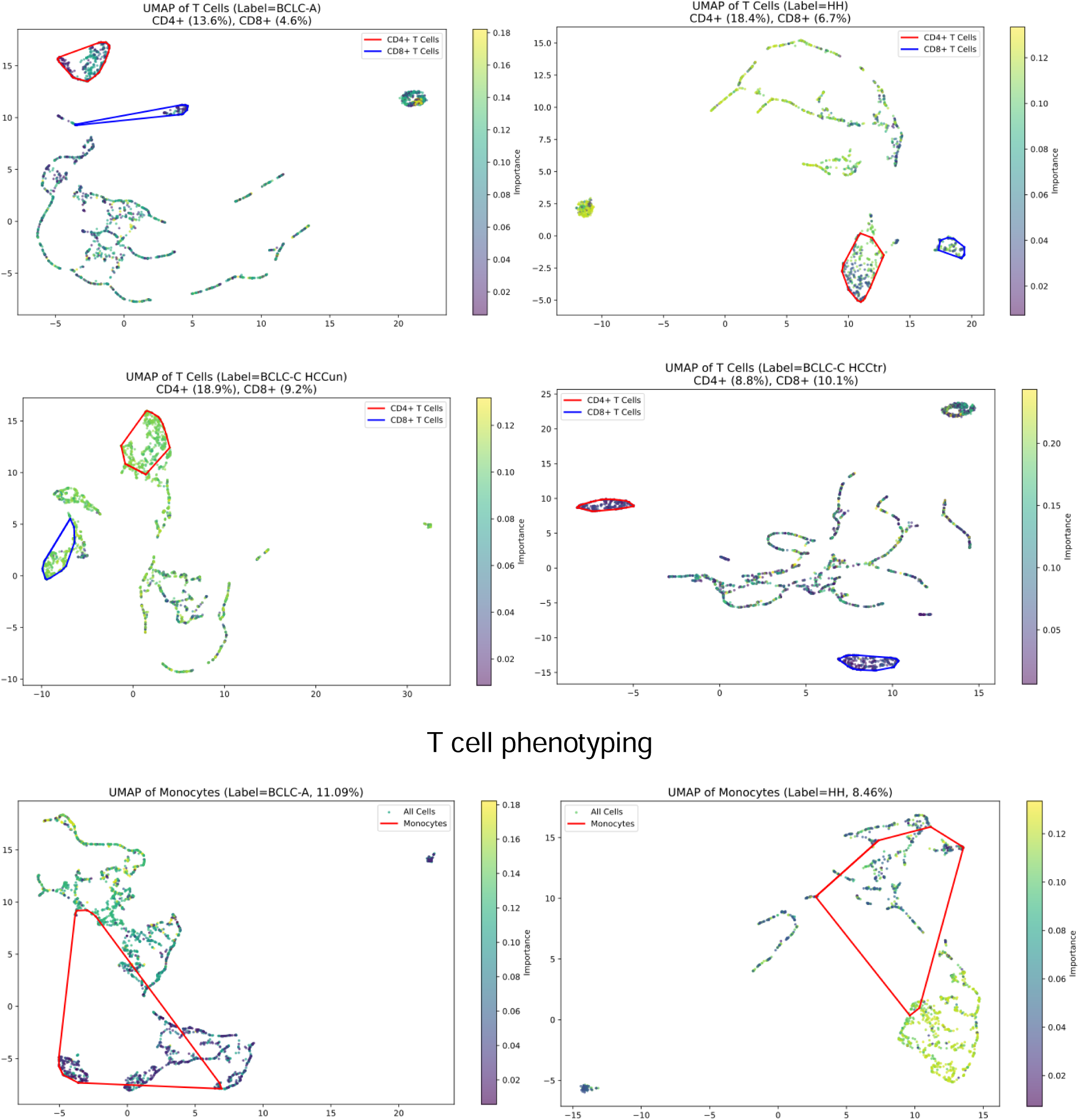
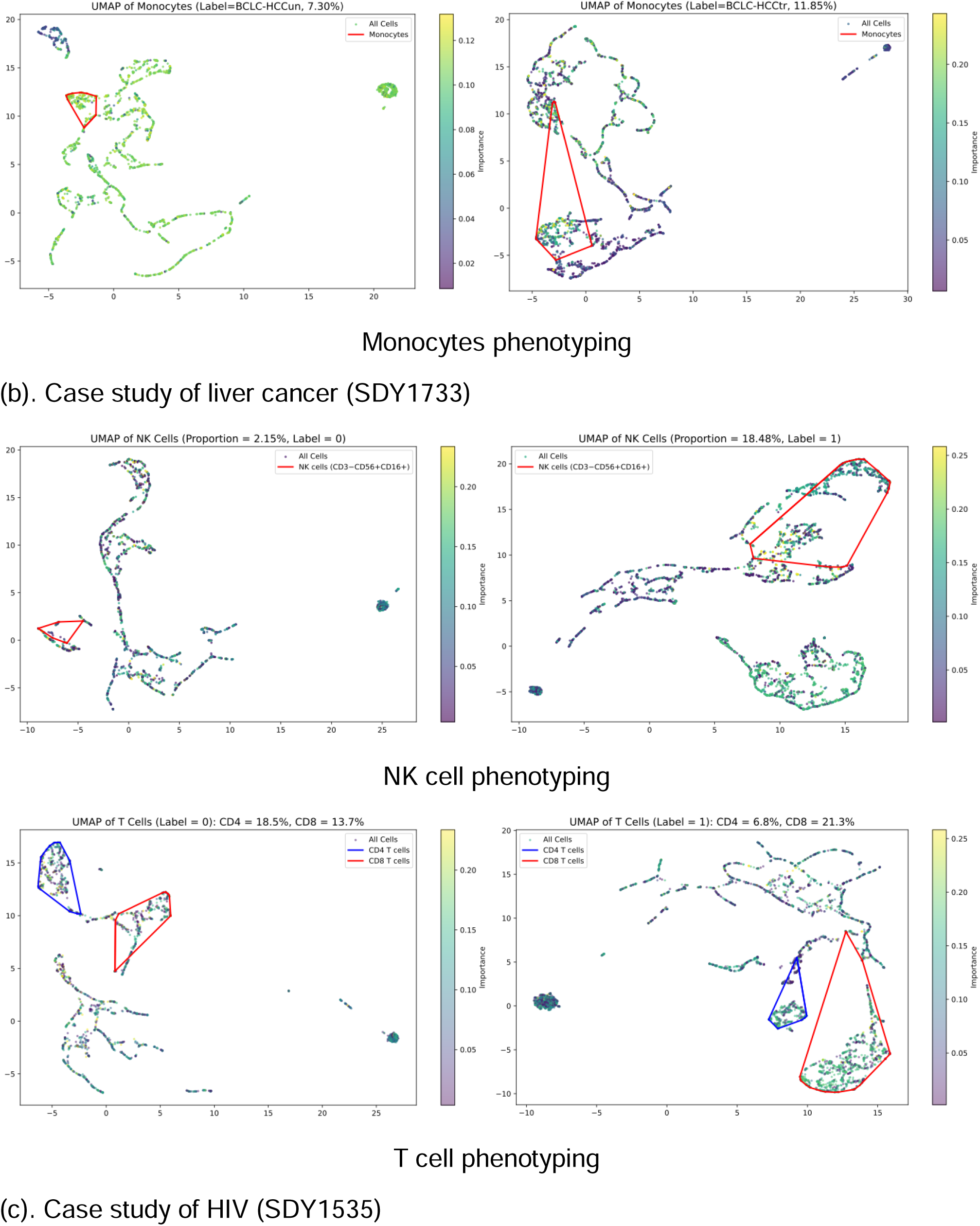
Bivariate plot of cell clusters. Each dot represents a single cell. The target cell types are identified by a red or blue circle. Color of the dot indicates importance.

#### Case study on COVID-19 (whole blood)

***SDY2011 data (COVID-19, human).*** This dataset is derived from a COVID-19 study^27^. The task is a binary classification of COVID-19 positive and negative. The samples are collected from serum, whole blood, and PBMC. There are 185 labeled samples. The data is available at SDY2011 on ImmPort.

**Important marker interpretation.** As shown in Figure 5 (a), the top 20 important markers are presented with importance calculated from the model and their expression in COVID-positive and COVID-negative groups. It reveals key patterns in the human immune response to SARS-CoV-2 infection. The prominence of myeloid markers (CD11b, CD11c, CD14) addresses the importance of monocyte activation in COVID-19 pathogenesis^39^. Notably, CD16 and CD66B show key patterns that potentially indicate neutrophil activity^40^. The prominence of CD39 suggests profound immunoregulatory alterations, possibly through adenosine-mediated suppression of effector functions^41^. CD45 maintains consistent importance as a pan-leukocyte activation marker that is important in both negative and positive groups. The high importance of CD64 points to Fcγ receptor-mediated mechanisms potentially leading to antibody-dependent enhancement. The pattern of CD123 possibly reflects plasmacytoid dendritic cell dysfunction, which has been validated in the previous study^42^.

From the expression difference as shown in Figure 5(a), notably, the CD39 and CD64 expression in the negative groups is significantly lower. CD39’s significantly lower expression in negative groups highlights its disease-specific role in COVID-19^41^. This clear expression pattern makes it a strong candidate for diagnosis and prognosis monitoring, especially for identifying immune exhaustion in severe cases. CD64’s minimal expression in negative samples also establishes it as a highly specific COVID-19 infection biomarker. Its significant upregulation indicates the profound monocyte/macrophage activation^43^.

**Important cell type interpretation.** Based on the important markers identified by ImmuneFM, we identified the important cell types based on these markers. Here we present and analyze a representative and important cell type, neutrophil cells, as shown in Figure 6(a). Our study utilized a combination of neutrophil-specific markers (CD16, CD66B, CD11B, and CD14) to identify the neutrophil and analyze its role in COVID-19. The results demonstrate a significant increase in neutrophil levels in positive groups, from a cell proportion of 27% to 46%. Neutrophils are among the first responders to viral infections^44^, consistent with their established role in early immune defense mechanisms.

The observed neutrophil elevation suggests their significant involvement in the host response to SARS-CoV-2 infection, which has been validated^27^. This quantitative change in neutrophil prevalence underscores their importance in COVID-19 pathogenesis and worth further investigation into their specific functional states during infection.

**Clinical and scientific insights.** Based on the analysis at the marker and cell levels, we summarized the key insights from clinical and scientific perspectives as follows.

1. **CD39 expression is significantly higher in COVID-19 patients, which could be a promising biomarker.** This point has been validated by several previous scientific studies. In the work of GB da Silva et al.^45^, they observed the increased expression of CD39 in moderate and severe cases, which has been stated as one of the key messages from their work. In the work of E Diaz-Garcia et al.^41^, they aim to quantify the relation between CD39 expression and severity in COVID-19 patients. They found that CD39 could be a promising biomarker for COVID-19 severity, and the overexpression of CD39 might be the reason for the disorder of thromboinflammation.
2. **CD64 could be an important COVID-19 biomarker with higher expression in positive patients.** This insight has also been validated by previous literature. In the study by M. Karawajczyk et al.^43^, CD64 is an early marker for COVID-19 infection and an indicator of severe cases. In the work of P. Bourgoin et al.^46^, they found that CD64 can be used to classify bacterial infection, COVID-19 infection, or other viral infections. This finding has the potential clinical application in the emergency department.
3. **Neutrophil cell increases in COVID-19 patients and play an important role in the immune response.** This point is also validated in the corresponding literature of SDY2011^27^. In the work by E McKenna et al.^47^, severe COVID-19 patients have an elevated level of neutrophil cells, which is an unusual situation in other types of viral infection. Neutrophils are also highly related to complications of COVID-19, e.g., thrombosis. In the review by LHA Cavalcante-Silva^48^, there are lots of previous efforts to investigate the neutrophil and COVID-19. The highlight is that COVID-19 triggers the activation of neutrophils.

#### Case study on live cancer staging (Whole blood)

***SDY1733 data (Oncology, human).*** This dataset is derived from a liver cancer stage research study^23^. The task is a multi-class classification of the liver cancer stage. The samples are collected from peripheral blood mononuclear cells (PBMC). We derived three live cancer stages, which are hepatic hemangioma (HH), stage A, stage C- untreated, and stage C-treated, based on the Barcelona Clinic Liver Cancer (BCLC) criteria. There are 100 labeled samples. The data is available at SDY1733 on ImmPort.

**Important marker interpretation.** As shown in Figure 5(b), we list the top 20 important markers and their expression difference in different HCC groups. We can find that the salient markers play important roles in both adaptive and innate immunity. T-cell regulators (CD3, TCR, CD8A, CD28, ICOS) indicated critical information about the T-cell activation and infiltration across different HCC stages. Particularly noteworthy, ICOS is essential in T-cell stimulation and immunotherapy^49^. B-cell markers like CD19 and IGD contribute to humoral immune responses. CD163 and CD172A/B are macrophage-associated molecules, providing insights into tumor-associated macrophages^50^. The identification of immune checkpoints (PD-1, TIM-3) was very important, as they not only indicated the T-cell exhaustion but also provided a target for therapy, e.g., anti-PD-1 immunotherapy. Proliferation and differentiation markers (KI67, TBET, CD127) provided valuable insights about the change of cell populations in HCC prognosis^51^. Notably, CD103 (tissue-resident memory T cells) and CCR7 (lymphocyte homing) suggested distinct stage-specific immune patterns. CD45 (pan-leukocyte markers) and CD11 (integrin family members) are related to the changes of leukocyte composition and cell adhesion dynamics during HCC prognosis.

**Important cell type interpretation.** As shown in Figure 6(b), we have several observations about the T cell phenotypes. Our analysis revealed distinct CD4+ and CD8+ T cell distribution patterns at different HCC stages. CD4+ T cells were significantly reduced in BCLC-C-treated groups, while CD8+ T cells showed lower proportions in BCLC-A HCC. Notably, total T cells (CD4+ and CD8+) were consistently diminished in both BCLC-A and BCLC-C treated groups compared to other stages, which has also been validated by the study of Shi et al.^23^, suggesting similar immunosuppressive microenvironments in these cohorts. This shared reduction in T cell infiltration may reflect progressive immune dysfunction or treatment-induced modulation, highlighting potential commonalities in immune evasion mechanisms between BCLC-A and treated advanced (HCCtr) HCC.

In addition to T cell alterations, we observed elevated monocyte proportions in both BCLC-A and BCLC-C treated groups compared to other HCC stages. This high-level monocyte indicates a potential shift toward myeloid-driven immunosuppression in these cohorts, which aligns with their shared T cell depletion phenotype that has also been validated in the study of Shi et al.^23^. The concurrent rise in monocytes possibly contribute to the tumor-permissive microenvironment in these stages, either through T cell suppression or tumor-associated macrophages polarization.

**Clinical and scientific insights.** Based on the marker and cell levels analysis, we summarized the key insights from clinical and scientific aspects as follows

1. **CD172a/b is an important marker in HCC staging classification**. From the expression analysis, CD172a/b has significantly higher expression only in the BCLC-C untreated stage. CD172a is an immune suppression receptor that inhibits the phagocytosis of tumor-associated macrophages. This point has also been validated in the previous research on the effectiveness of 3-HAA to HCC^52^.
2. **Decreasing the ICOS expression level might be an effective immunotherapy approach for HCC.** There is previous work that validates this point. In the study conducted by Lu et al^53^, higher level ICOS+ Tregs are related to worse overall survival. Depleting the ICOS Tregs may be an effective way to improve the clinical outcomes of HCC patients. Because the original study^23^ with this data used anti-PD-1 immunotherapy as treatment, this data-driven finding about ICOS may implicitly indicate the mechanism of anti-PD-1 and ICOS expression inhibition.
3. **T cells are increased and monocytes are decreased in the BCLC-A groups and the BCLC-C treated groups of HCC.** This has been validated by the study that generated this dataset^23^. This finding further underscores the distinct immune landscape of BCLC-A and treated BCLC-C stages. It implicates the myeloid cells as potential and promising therapeutic targets or biomarkers in the HCC prognosis.

#### Case study on HIV diagnosis (PBMC)

**Important marker interpretation.** In Figure 5(c), the top 20 important markers identified by ImmuneFM are shown. CD4 and CD19 represent central T helper and B cell compartments, always reduced and impaired by HIV. Notably, PD-1 and KI67 reflect T cell proliferation and exhaustion, which can be used as landscapes of chronic HIC infection^54^. HLA-DR, CD54, and CD123 indicate activation of myeloid and dendritic cells. Innate immunity also emerges as essential, with NKG2A, NKp44, KIR2DS4, and ULBP2-5-6 implicating NK cell activation^55^. The presence of HLA-E and HLA-G further indicates changes in non-classical antigen presentation. Markers CD304 and CD112 and chemokine CXCL13 indicate dysregulated lymphoid and immune cell communication^56^. FCRγ is related to antibody-mediated cytotoxicity and phagocytosis, which can be impaired by HIV. Moreover, plasmacytoid dendritic cell markers like CD303 suggest altered interferon responses.

As shown in the marker expression difference in Figure 5(c), the HIV-positive group exhibits significantly lower expression of CD19, CD4, CD54, and CD7, indicating impaired B and T cell function as well as reduced immune adhesion and signaling^57^. On the other hand, increased expression of FCRγ and KI67 reflects heightened immune activation and cell proliferation in response to HIV infection^58^. These expression differences highlight the multifaceted immune dysregulation associated with HIV.

**Important cell type interpretation.** As shown in Figure 6(c), Natural Killer (NK) cells increased in the HIV-positive groups. This has also been validated in the study of this dataset by Vendrame et al.^26^. This elevation often comes with phenotypic and functional changes, e.g., changed expression of activating and inhibitory receptors and reduced cytotoxicity. Although the proportion is elevated, these NK cells may show signs of exhaustion or dysfunction, which limits their effectiveness and capability to target HIV-infected cells. The alteration of NK cells may be related to immune evasion and provide a promising target for HIV immunotherapy.

Additionally, we also observed CD4 T cells decreasing and CD8 T cells increasing in the HIV positive groups. CD4 T cell is the primary target of HIV, causing impaired adaptive immunity. The CD4 loss is accompanied by a relative expansion of CD8 T cells that proliferate in response to antigen exposure. Although the CD8 T cell proportion is elevated, they are often exhausted with markers such as PD-1, TIGIT^26^. The imbalance between CD4 and CD8 T cells reflects ongoing immune dysregulation and activation. The total number of CD4 and CD8 cells has also decreased. Monitoring these alterations provides essential insights into HIV prognosis and potential immunotherapy strategies.

**Clinical and scientific insights.** Based on the marker and cell-level interpretation analysis, we summarize the clinical and scientific insights as follows.

1. **CD7 decreases in the HIV infected patients.** This point has been validated by previous scientific evidence. Aandahl et al^59^. found that the loss of high-expression CD7 cells is related to HIV infection, especially in patients with fast progression. The reduced CD7 level is caused by the expansion of CD8 T cells with low CD7 expression.
2. **KI67 increases in the HIV infected patients.** This insight was also validated in previous literature. Sachsenberg et al.^60^ measured CD4 and CD8 T cell turnover in HIV patients by KI67. This was also found in our phenotyping results of T cells in Figure 6(c). The KI67 indicating CD4/CD8 T cells turnover provides promising direction for immunotherapy of HIV.
3. **NK cells and CD8 T cells increase, but CD4 T cell decreases in the HIV infected patients.** This point was validated in the original study^26^ of this HIV downstream data. TIGIT-marked NK cells were elevated in the HIV infected groups. CD4 T cells are known to be the target of HIV, which will significantly decrease after infection. Elevated levels of CD8 T cells may indicate an exhaustion of it. The phenotyping of these immune cells provides promising insights into possible treatment for HIV^61^.

## Discussion

In this work, we curated an AI-ready large-scale cytometry dataset from the ImmPort platform. It is a dataset covering a wide range of immunology-related diseases and conditions. Due to the affordable expense and widespread usage, cytometry data is the most common assay type in immunology. However, various independent studies have different goals and aims. Manual efforts are needed to integrate multi-source datasets. This dataset-level contribution aims to address the lack of a large-scale and comprehensive immunology dataset. It can be used to pre-train the immunology-related foundation model and fine-tune other foundation models. Through building this dataset, 203 commonly used panel markers are included, which can cover most cytometry results analysis.

With the Transformer used as the backbone of ImmuneFM, the model with the attention mechanism is able to capture the relation between markers and learn the embeddings of both markers and cells simultaneously. Through the self-supervised learning on a large-scale cytometry dataset, ImmuneFM gains the generalizability across different immunology fields. Compared to the plain Transformer without pre-training, the pre-trained ImmuneFM achieves significantly better performance, which implies the necessity of pre-training on large-scale data. The pre-trained ImmuneFM demonstrates superior performance with a limited number of labeled samples, which is a common challenging scenario in biomedicine.

From the quantitative performance results, ImmuneFM outperforms the CNN, Transformer trained from scratch. Moreover, ImmuneFM also outperforms the traditional method based on feature engineering, e.g., FlowSOM+GBDT. It demonstrates the effectiveness of pre-training on large-scale data. Supervised model, including CNN and Transformer, needs enough labeled samples for training, so their performance is not satisfactory with a limited number of labeled cytometry samples. Moreover, the traditional method of feature engineering demonstrates good performance and outperforms supervised deep learning models on some tasks. The feature engineering method, like FlowSOM, can extract the cell distribution information with a limited number of samples, which contributes to the competitive performance.

ImmuneFM also accelerates the clinical and scientific discovery in a data-driven way. We analyze three case studies including cancer, HIV, and COVID-19. Through the classification of cytometry samples and the interpretability of ImmuneFM, we can identify the important biomarkers and single cells that contribute to the sample diagnosis. This data-driven method can identify scientific discoveries which has been validated by previous literature and evidence from the wet lab. The immunology foundation model is a good example of how we could use AI for scientific discovery. In the future, we plan to extend this framework to multimodal immunology data, including multi-omics, tabular data, images, and text. Additionally, supporting more immunology-related analytic tasks is a promising direction.

